# Mammary-specific ectopic expression of mutant human GATA3 did not potentiate medroxyprogesterone acetate-driven mouse mammary tumorigenesis

**DOI:** 10.1101/2023.01.03.522644

**Authors:** Kristopher A. Lofgren, Paraic A. Kenny

## Abstract

GATA3 is somatically mutated in approximately 15% of estrogen receptor positive human breast tumors, however the mechanism(s) by which these alterations contribute to tumorigenesis are unclear. The observed patterns of mutations suggest a strong selective pressure to mutate a single allele of GATA3 in a manner favoring retention of the first of two zinc finger domains. The non-mutated GATA3 allele is maintained and expressed. We and others have hypothesized that expression of the mutant GATA3 protein may actively contribute to breast tumorigenesis, however the aging of several independently generated mouse models with mammary-specific mutant GATA3 expression did not result in tumorigenesis. In this study, we evaluated whether a mammary tumor-promoting dose of medroxyprogesterone acetate could synergize with mammary specific mutant GATA3 (G335fs) expression and accelerate the kinetics of tumor formation. We report that the tumor incidence rate in these animals did not differ from that observed in wild-type littermate controls.

## INTRODUCTION

GATA3, a transcription factor expressed in luminal mammary epithelial cells, is a key regulator of mammary gland development (1). In breast cancer, GATA3 is expressed in estrogen receptor positive (ER+) tumors (2,3) where GATA3 and ERα are reciprocally positively regulated (4). Somatic GATA3 mutations occur in approximately 15% of ER+ tumors (5-7) but the mechanism(s) by which these alterations contribute to tumorigenesis remain unclear.

GATA3 mutations in breast cancer are clustered in exons 4, 5 and 6 which encode the zinc finger domains necessary for DNA binding. In addition, the wild-type GATA3 allele tends to be retained and both alleles are expressed in these tumors (7,8). Approximately 40% of the mutations identified in the TCGA study lack the second zinc finger, which is responsible for binding the canonical GATA motif (9,10), while the remainder encode longer proteins containing both of the GATA3 zinc fingers. Initially, it was proposed that GATA3 was a haploinsufficient tumor suppressor gene (7), however recent studies indicate that the mutant proteins encoded by these alleles may have pro-tumor properties, evidenced by promotion of the growth of xenografted human breast cancer cells (11-13).

We previously described a transgenic mouse model in which human mutant GATA3 (GATA3^335fs^) is expressed in the mammary gland under the control of the MMTV enhancer, resulting in precocious lobuloalveolar development but no mammary tumor formation in aged cohorts of female mice (11). This mutation is a recurrent hotspot in human breast cancer. More recently, another group using a different genetic engineering approach to generate mouse lines ectopically expressing two different truncating mouse GATA3 mutant alleles (both corresponding to hotspots in human breast cancer) did not detect an impact on mammary gland architecture but, as with our study, also did not detect tumor formation in aged mice (14).

Together, these two studies (11,14) demonstrate that ectopic expression of mutant GATA3 in animals bearing two wild-type GATA3 alleles is insufficient to promote mammary tumor development. This leaves unresolved competing hypothesis about whether either an intrinsic gain-of-function activity of truncating GATA3 mutations, or GATA3 haplo-insufficiency, or a combination might be providing the selective pressure that results in GATA3 being among the most frequently mutated genes in estrogen-receptor positive breast cancer. Because models with rapid tumor onset can mask the contribution of a potentially weak modifier, we believed a long latency model was optimal to test the hypothesis that expression of mutant GATA3 could accelerate the kinetics of mammary tumor formation. Implantation of slow-release medroxyprogesterone acetate (MPA) reliably induces mouse mammary tumors with relatively long latency (52 weeks), 80% of which express the estrogen and progesterone receptors (15-17). As a step toward resolving this issue, we implanted female MMTV-GATA3^335fs^ mice and wild-type littermates with MPA and monitored tumor development.

## MATERIALS & METHODS

### Mouse studies

All experimentation was approved by the Institutional Animal Care and Utilization Committee of the University of Wisconsin, La Crosse. The generation and characterization of FVB mice ectopically expressing human mutant GATA3 under the control of the Mouse Mammary Tumor Virus Long Terminal Repeat, FVB-Tg(MMTV-GATA3^G335fs^)1Kny, was previously described (11) and are simply referred to as MMTV-GATA3^G335fs^ hereafter. Control mice were transgene-negative female FVB littermates. Female mice were injected subcutaneously with a slow-release MPA formulation in the dorsal scruff between 6 and 9 weeks of age. Mice were monitored weekly for mammary tumorigenesis. Date of first palpable lesion was recorded in each case. All mice were subjected to necropsy and, if a previously undetected mammary tumor was noted on necropsy, the date of euthanasia was used as the date of tumor detection.

### Slow-release medroxyprogesterone acetate (MPA)

Medroxyprogesterone acetate was compounded at 160 mg/mL for subcutaneous injection with USP grade materials by a local compounding pharmacy (The Prescription Center, La Crosse, WI). The formulation included methylparaben, propylparaben, polyethylene glycol, polysorbate 80, methionine, povidone, sodium chloride, monobasic and dibasic sodium phosphate, and water as inactive ingredients. The pH was adjusted to 6.9 with sodium hydroxide. Potency, stability, sterility, and endotoxin testing was performed by Eagle Analytical Services (Houston, TX). Potency test results averaged 94%, thus 90 mg of MPA was injected to keep the volume being administered to an acceptable scale, relative to body size. Efficacy of dosing was confirmed by estrus checks that illustrated cessation of estrous cycling.

### Statistics

All statistical analyses were performed using GraphPad Prism (ver. 9.3.1; Graphpad Software, La Jolla, CA, USA). Survival analysis was by the method of Kaplan and Meier (18). Statistical significance was evaluated using the log-rank test with 0.05 as the significance threshold.

## RESULTS

To provide a moderate pro-tumorigenic stimulus, 21 wild-type and 25 GATA3^335fs^ mice were implanted with a single bolus of slow-release MPA between ages 6 and 9 weeks and monitored for tumor formation. Median follow-up times for the wild-type and MMTV-GATA3^335fs^ mice were 525 and 522 days, respectively. The earliest tumor was detected 341 days post MPA injection. In total, 28% of the wild-type mice and 20% of the MMTV-GATA3^335fs^ mice developed mammary tumors during follow-up (χ^2^ p-value = 0.497). Kaplan-Meier analysis of time to tumor onset (Fig 1) showed no difference between the two genotypes.

**Figure 1.**
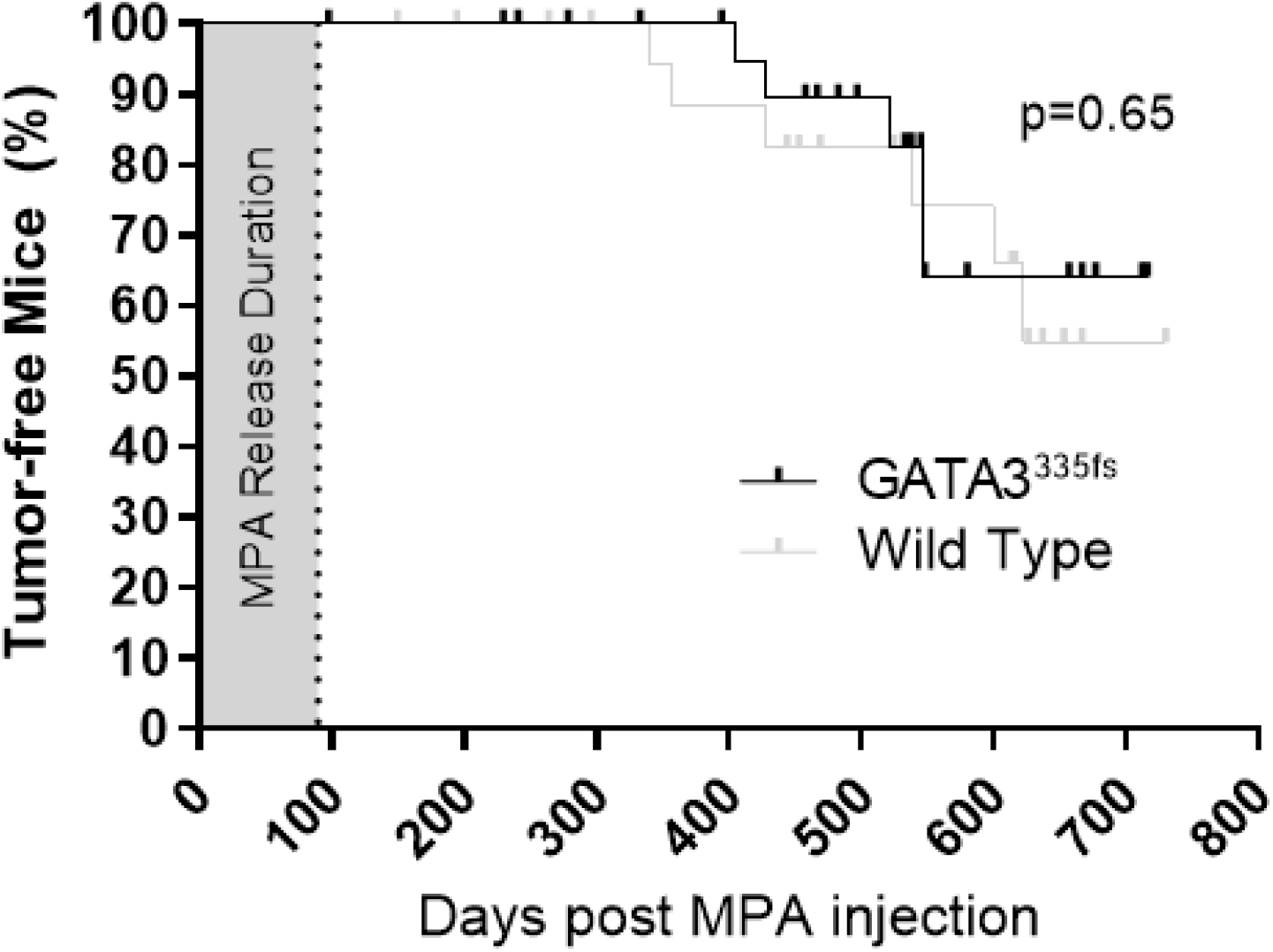
Ectopic expression of GATA3^335fs^ does not alter the rate of medroxyprogesterone acetate induced mammary tumorigenesis. Kaplan-Meier analysis demonstrating time to tumor detection in cohorts of female MMTV-GATA3^335fs^ mice (n = 21) or their wild-type littermates (n = 25). The approximate duration of MPA exposure from the single bolus of slow-release gel is indicated by shading.

## DISCUSSION

Despite the prevalence of GATA3 mutations in ER+ human breast cancer, the mechanism(s) by which these genetic alterations contribute to breast tumor initiation and/or progression are unresolved. Even whether GATA3 is an oncogene, a tumor-suppressor gene or shares some properties of both categories in a context-dependent manner remains unclear. Following up on two prior independent studies (11,14) that showed that ectopic expression of various hotspot GATA3 mutations failed to elicit mouse mammary tumor formation, we asked here whether the combination of mutant GATA3 expression with a modestly potent mammary tumor initiating treatment could synergize to promote faster tumorigenesis. We conclude that mammary-specific expression of human GATA3^335fs^ mutant did not potentiate MPA-driven tumorigenesis in mice.

Although the study was negative, it does represent an incremental advance in addressing the overarching question of the mechanism(s) by which GATA3 alterations contribute to breast tumorigenesis. Fundamentally, we need to understand whether the effect of the heterozygous GATA3 mutations prevalent in breast cancer represent a loss-of-function tumor suppressor phenomenon or are more consistent with an active mechanistic contribution of the mutant proteins to some tumor-relevant process, in a manner typically consistent with the definition of an oncogene.

Evidence in favor of an oncogene-like role includes (a) the introduction of mutant GATA3 enhances xenograft tumor growth by both ZR-75-1 (11) and CAMA-1 (12) breast cancer cell lines, (b) the ability of mutant GATA3 proteins to tether to DNA and influence gene transcription (13,14,19), (c) the demonstration that repairing the GATA3 mutation in MCF7 cells using recombination can attenuate tumor growth (12) and (d) the observed pattern of mutation accumulation (11,12,14,20) in hotspots (typical of oncogenes) as opposed to a more diverse distribution of stop or frame-shift mutations throughout the coding sequence, or even larger deletions, which would be more typical of a tumor suppressor gene. Data on the impact of ectopic expression of mutant GATA3 on mammary architecture are mixed, with one study finding no effect (14) and another reporting precocious lobuloalveolar development (11). A strength of both studies was the long-term follow-up on these animals which demonstrated no increased mammary tumorigenesis. The present study showed that ectopic mutant GATA3 expression did not synergize with a tumor promoting hormonal stimulation model to increase the rate of tumorigenesis, a finding that does not support the oncogene hypothesis.

Despite this lack of clarity regarding an oncogene-like role, the hypothesis that GATA3 functions like a classical two-hit tumor suppressor gene has little evidentiary support. As already stated, the observed pattern of mutations, which implies a strong selective pressure to maintain expression of a truncated GATA3 protein that includes the first zinc finger domain, is not consistent with simple elimination of GATA3 conferring a tumor-relevant advantage. In human ER+ breast tumors with GATA3 mutations, one wild-type allele is retained and expressed indicating that, whatever the mechanism of action is, it does not depend on elimination of functional GATA3 from the cell lineage. If GATA3 is not a classical two-hit tumor suppressor gene, could a single copy mutation resulting in haploinsufficiency be the underlying mechanism? Again, the specificity of observed pattern of mutations strongly suggests that elimination of the function of one allele does not underlie the pro-tumorigenic effect. GATA3 expression is a defining feature of luminal ER+ breast cancer and much work has demonstrated that re-introduction of GATA3 expression in GATA3-negative triple-negative breast cancer cell lines can attenuate tumor growth in xenografts (21,22). However, we believe that these experiments are more relevant to consideration of the impact of GATA3-controlled transcriptional programs on differentiation rather than supporting the contention that GATA3 is a classical tumor-suppressor gene in the absence of any cancer genetics evidence associating GATA3 mutational loss with the initiation or progression of ER-negative breast cancer.

Lying between these two competing hypotheses is a hybrid which has not yet been investigated experimentally. Clearly, in human breast cancer, the dose of wild-type GATA3 is reduced by half, while the tumor lineage gains a mutant GATA3 allele. Both our mouse model (11) and the one from the Jonkers group (14) rely on ectopic expression of mutant GATA3 in the context of a mouse with two intact wild-type GATA3 alleles, an experimental design that is generally adequate to detect a pro-tumor signal from potent oncogenes like ERBB2 (23). If the net pro-tumor effect of the recurrent GATA3 mutations is to simultaneously reduce the GATA3 dose by half and to express a truncated GATA3 with some residual or neo-morphic pro-tumorigenic function, then experimentation using a floxed GATA3 or otherwise engineered model that more accurately recapitulates the gene dosage reduction in human tumors will be required to resolve these important outstanding issues. Additionally, cell culture experiments that do not simply repair the GATA3 mutation (12) but instead selectively eliminate it may permit a more nuanced separation of oncogene and haploinsufficiency interpretations.

## ACKNOWLEDGEMENTS

This work was supported by the Gundersen Medical Foundation. PK holds the Dr. Jon & Betty Kabara Endowed Chair in Precision Oncology. KL was supported, in part, by the Norman L. Gillette Jr Breast Cancer Research Fellowship.

